# Environmentally-relevant doses of bisphenol A and S exposure in utero disrupt germ cell programming across generations resolved by single nucleus multi-omics

**DOI:** 10.1101/2024.12.05.627072

**Authors:** Liang Zhao, Mingxin Shi, Sarayut Winuthayanon, James A. MacLean, Kanako Hayashi

## Abstract

**Background:** Exposure to endocrine-disrupting chemicals (EDCs), such as bisphenol A (BPA), disrupts reproduction across generations. Germ cell epigenetic alterations are proposed to bridge transgenerational reproductive defects resulting from EDCs. Previously, we have shown that prenatal exposure to environmentally relevant doses of BPA or its substitute, BPS, caused transgenerationally maintained reproductive impairments associated with neonatal spermatogonial epigenetic changes in male mice. While epigenetic alterations in germ cells can lead to transgenerational phenotypic variations, the mechanisms sustaining these changes across generations remain unclear.

**Objectives:** This study aimed to systematically elucidate the mechanism of transgenerational inherence by prenatal BPA and BPS exposure in the murine germline from F1 to F3 generations at both transcriptomic and epigenetic levels.

**Methods:** BPA or BPS with doses of 0 (vehicle control), 0.5, 50, or 1000 µg/kg/b.w./day was orally administered to pregnant CD-1 females (F0) from gestational day 7 to birth. Sperm counts and motility were examined in F1, F2, and F3 adult males. THY1^+^ germ cells on postnatal day 6 from F1, F2, and F3 males at a dose of 50 µg/kg/b.w./day were used for analysis by single-nucleus (sn) multi-omics (paired snRNA-seq and snATAC-seq on the same nucleus).

**Results:** Prenatal exposure to BPA and BPS with 0.5, 50, and 1000 µg/kg/b.w./day reduced sperm counts in mice across F1 to F3 generations. In the F1 neonatal germ cells, ancestral BPA or BPS exposure with 50 µg/kg/b.w./day resulted in increased differentially expressed genes (DEGs) associated with spermatogonial differentiation. It also disrupted the balance between maintaining the undifferentiated and differentiating spermatogonial populations. Differentially accessible peaks (DAPs) by snATAC-seq were primarily located in the promoter regions, with elevated activity of key transcription factors, including SP1, SP4, and DMRT1. Throughout F1-F3 generations, biological processes related to mitosis/meiosis and metabolic pathways were substantially up-regulated in BPA-or BPS-exposed groups. While the quantities of DEGs and DAPs were similar in F1 and F2 spermatogonia, with both showing a significant reduction in F3. Notably, approximately 80% of DAPs in F1 and F2 spermatogonia overlapped with histone post-translational modifications linked to transcription activation, such as H3K4me1/2/3 and H3K27ac. Although BPA exerted more potent effects on gene expression in F1 spermatogonia, BPS induced longer-lasting effects on spermatogonial differentiation across F1 to F3 males. Interestingly, DMRT1 motif activity was persistently elevated across all three generations following ancestral BPA or BPS exposure.

**Discussion:** Our work provides the first systematic analyses for understanding the transgenerational dynamics of gene expression and chromatin landscape following prenatal exposure to BPA or BPS in neonatal spermatogonia. These results suggest that prenatal exposure to environmentally relevant doses of BPA or BPS alters chromatin accessibility and transcription factor motif activities, consequently contributing to disrupted transcriptional levels in neonatal germ cells, and some are sustained to F3 generations, ultimately leading to the reduction of sperm counts in adults.

## 1. Introduction

Endocrine-disrupting chemicals (EDCs) are exogenous chemicals present in the environment that interfere with the endocrine system, altering normal homeostatic regulation and developmental processes^1^. Exposure to EDCs has been linked to various health issues, particularly reproductive and metabolic disorders^2–4^. Maternal EDC exposure is especially concerning, as fetal and neonatal stages are highly susceptible to hormonal and genetic disruptions^5,6^. Reproductive and developmental defects, including compromised fertility and adverse neurodevelopmental outcomes, have been associated with early-life EDC exposure^7–9^. Moreover, evidence suggests that these EDC-induced harmful effects can extend beyond the directly exposed generation and be transmitted to future generations^10–17^. These findings have raised significant public health concerns and an urgent need for a deeper understanding of the transgenerational inherence of EDC-dependent maladies.

Among EDCs, bisphenol A (BPA), a synthetic plasticizer widely used in manufacturing polycarbonate plastics and epoxy resins, is one of the most studied^18,19^. BPA exposure has been linked to a myriad of deleterious effects, including reproductive and developmental abnormalities, cancer, neurobehavioral disorders, metabolic syndromes, and cardiovascular diseases^20,21^. Consequently, BPA analogs, such as bisphenol S (BPS), were introduced as “safer” alternatives^22,23^. However, growing evidence has revealed that BPS, elicits toxic effects similar to BPA, especially on the reproductive system^22–24^. In males, epidemiological studies have shown a direct correlation between BPA or BPS exposure and lower sperm counts and motility^25–30^, abnormal sperm morphology^27,30,31^, and damaged sperm DNA integrity^30^. Rigorous prior studies, including from our group, indicate that exposure to BPA and BPS during critical periods of development resulted in reproductive abnormalities that persist for generations^32–38^. Furthermore, BPA and BPS were detected in breast milk as the predominant bisphenol compounds in Asia, America, and Europe^39,40^. Children and adolescents have even higher urinary BPA and BPS concentrations than adults^41,42^, heightening their health risks to future generations.

In mammals, male germ cell development involves a sequential progression of cell fate transitions^43^. In the embryo, primordial germ cells (PGCs) undergo mitotic proliferation and form prospermatogonia, which transit into undifferentiated spermatogonia after birth^44^. Neonatal undifferentiated spermatogonia consist of spermatogonial stem cells (SSCs) and spermatogonial progenitor cells^45,46^. SSCs maintain spermatogenesis through self-renewal and differentiation, while progenitor cells primarily commit to differentiation but retain the ability to regenerate self-renewing capacity^47,48^. Proper germ cell development and function requires precise epigenetic regulation at key stages, including genome-wide epigenetic reprogramming during prospermatogonia formation and epigenetic fine-tuning in SSCs^49,50^. However, exposure to EDCs can disrupt these processes, leading to heritable aberrant epigenetic marks, potentially driving transgenerational phenotypic alterations^5,51,52^. Rodent studies have demonstrated that BPA and BPS exposure alters epigenetic patterns in male germ cells, including changes in DNA methylation and histone modifications, such as H3K9me3 and H3K27me3^53,54,55,56,57^. Our group has recently reported that ancestral prenatal exposure (F0) to BPA and BPS transgenerationally induced lower sperm counts and motility in mice, accompanied by elevated DNMT3B and diminished H3K9me2 and H3K9me3 in the F3 neonatal spermatogonia^35^. Similarly, gestational exposure to a mixture of BPA and phthalates was found to promote epigenetic transgenerational inheritance of reproductive disease and sperm epimutations^58^. Rahman et al. observed that altered DNA methylation in adult spermatozoa of the F3 mice correlated with decreased sperm counts following gestational BPA exposure in the F0 pregnant females^32,34^. These findings highlight the critical impact of epigenetic disruptions caused by EDC exposure on germline integrity and male fertility across generations.

Given the complexity of cell fate transitions during germ cell development^43,46^, it is important to elucidate the specific changes of germ cell subpopulations affected by EDCs. Recently, an increasing knowledge of male germ cell development has been obtained using single-cell RNA sequencing (scRNA-seq) and single-cell sequencing assay for transposase-accessible chromatin (scATAC-seq)^43,45,46,59,60^. Moreover, single-cell multi-omics sequencing now enables the simultaneous profiling of transcriptomes integrated with its chromatin accessibility at the single-cell level. While some studies have used scRNA-seq to investigate the effects of EDC exposure on germ cell transcriptome, they have been limited by the utilization of a potentially toxic high dose (e.g. DEHP at 750 mg/kg body weight) or were conducted using in vitro exposures^61,62^, which may not accurately represent the daily physiological exposure paradigm. To date, there has been no rigorous investigation into EDC-dependent epigenetic changes in germ cells and their heritable mechanisms at single-cell resolution.

In this study, we focus on postnatal day 6 (PND6) spermatogonia to investigate the transgenerational impacts of prenatal exposure to environmentally relevant doses of BPA and BPS in mice. PND6 spermatogonia were chosen as they represent a developmental stage at which epigenetic reprogramming is nearly completed. Using integrated single-cell RNA and ATAC sequencing, we generated paired, germ cell-specific chromatin accessibility and transcriptional profiles from the same cell for deeper dissection of transgenerational impacts from BPA and BPS prenatal exposure with an environmental dose in mice. To our knowledge, this is the first study that leverages information from scRNA-seq and scATAC-seq to systemically unveil the transgenerational dynamics of gene expression and chromatin landscape on neonatal male germ cells. We identified numerous conserved and altered germline transcriptomic and chromatin features across generations. Specifically, prenatal BPA or BPS exposure disrupted the balance between maintaining the undifferentiated and differentiating spermatogonial populations in the F1 generation. BPA and BPS exposure mainly altered chromatin accessibility in the promotor regions of germ cells. Interestingly, DMRT1 motif activity was consistently elevated following ancestral exposure to BPA or BPS among three generations, which may drive the disturbance of germ cell homeostasis and decrease sperm counts in offspring transgenerationally. These findings offered new insights into the mechanisms underlying the transgenerational inheritance of EDC effects.

## 2. Methods

### 2.1. Chemicals

BPA (#239658) and BPS (#103039) were purchased from Sigma-Aldrich. Tocopherol-stripped corn oil (ICN90141584) was purchased from Fisher Scientific. BPA and BPS were first dissolved in 100% ethanol and then diluted in tocopherol-stripped corn oil as previously described^35,63^.

### 2.2. Animals

Adult CD1 mice were purchased from Inotiv. All experiments utilizing animals were maintained in the vivarium and approved by Washington State University according to institutional guidelines for the care and use of laboratory animals (protocol #6751).

### 2.3. Study design

To investigate the transgenerational impacts of prenatal BPA and BPS exposure on the male germline, we devised an experimental strategy to allow the paternal transmission of the exposure effects (Figure 1a). Pregnant CD1 females (F0) were orally administrated with vehicle control (tocopherol-stripped corn oil), 0.5, 50, or 1000 µg/kg body weight per day (b.w./day) doses of either BPA or BPS (n=5-9 each group) from gestational day 7 (GD7, GD1 was defined as the presence of a vaginal plug) to birth. Daily oral feeding of BPA or BPS was performed by pipetting the tocopherol-stripped corn oil containing the dose into the mouth for better mimicking human BP intake as described previously^35,63^. We chose the dose range as the FDA has determined that no observed adverse effect level (NOAEL) for BPA is 5 mg/kg/b.w./day^64^. BPA dietary intake has been estimated at 0.5 µg/kg/b.w./day^65^. The body weight gain of dams was measured once a day to adjust the dosing. Mice delivered from the F0 females were labeled as F1 generation. At 6-7 weeks of age, male mice in the F1 generation were used to breed with untreated CD1 females to generate the F2 generation. With the same strategy, F3 generation was generated from F2 males.

**Figure 1.**
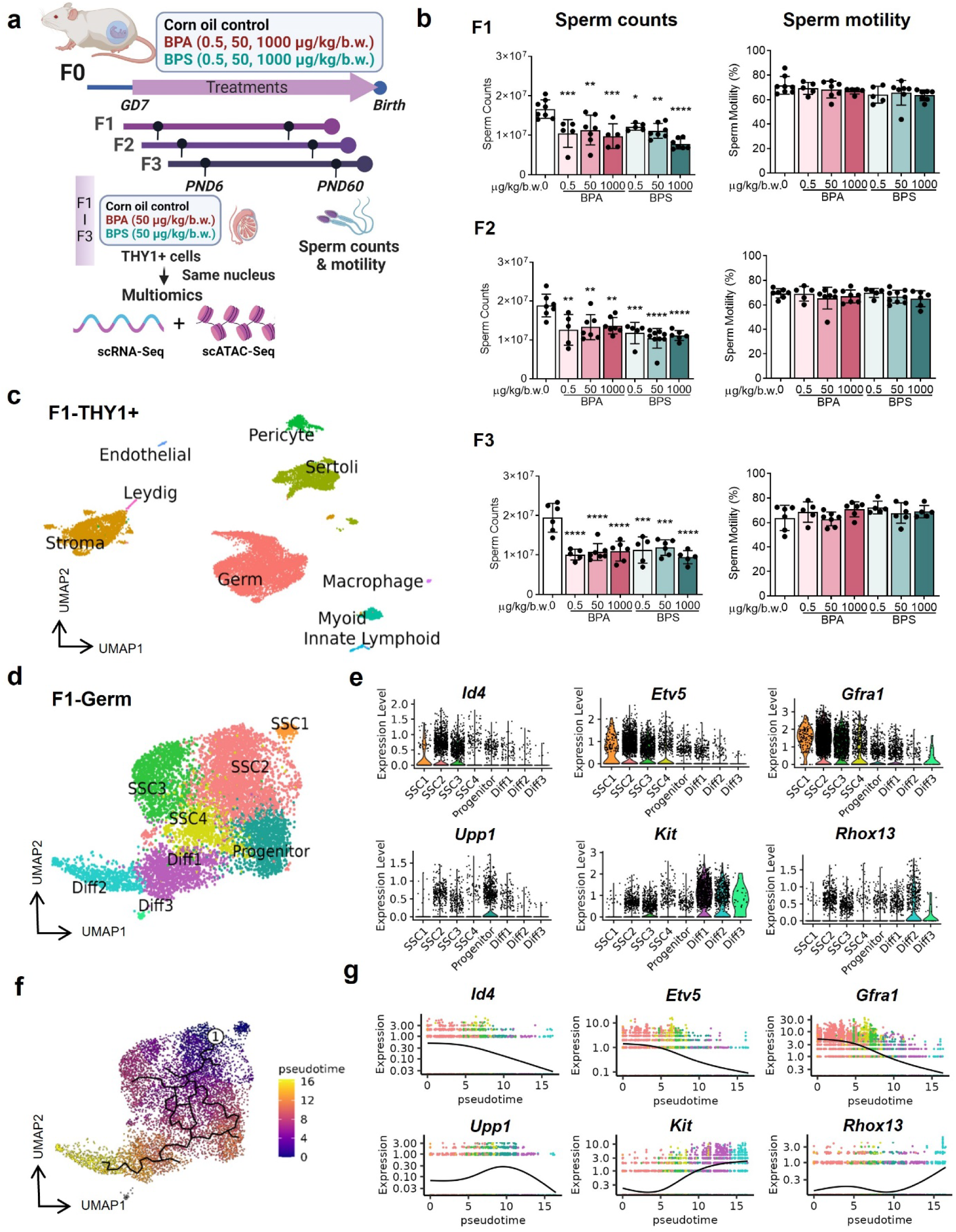
Experimental design and classification of THY1^+^ testicular cells in the F1 generation. (a) Schematic diagram of the experimental study design. Created with BioRender.com. (b) Sperm counts (left panel) and motility (right panel) across F1 to F3 generations. (c) WNN UMAP visualization of nine major cell types from PND6 testes in the F1 generation. (d) UMAP plot of germ cell subsets defined by clustering analysis. (e) Violin plots of representative marker genes for germ cell subtypes. (f) Monocle pseudotime trajectory analysis of the germ cell subsets defined in (d). Black lines on the UMAP represent the trajectory graph. The root is labeled with a circled 1. (g) Plots showing the expression pattern of representative germ cell marker genes along the pseudotime axis. WNN, “weighted-nearest neighbor” analysis.

On PND60, 1-2 male littermates (n=5-9/group/generation. The average from the same litter mates was used for n=1.) were euthanized, and body and paired testis weights were recorded. Both caudal epididymides were collected to assess sperm counts and motility. For multiome analysis, on PND6, 1-2 male littermates from at least 3 litters in each group were euthanized to collect neonatal testis. The dosage of 50 µg/kg/b.w./day was selected for the multiome analysis, as it was proposed as a chronic reference dose of oral BPA exposure in humans by the Environmental Protection Agency (EPA)^66^. Moreover, we have demonstrated that prenatal exposure to BPA and BPS at 50 µg/kg/b.w./day caused transgenerational reproductive defects, including lower sperm counts, in F3 males associated with altered expression of DNA methyltransferases and histone marks in the F3 neonatal testes^35^.

### 2.2. Sperm counts and motility

Sperm counts and motility were examined following our prior studies^35,63,67^. Briefly, two caudal epididymis were dissected and placed in 1 mL of EmbryoMax heated to 37°C. After 15 min of incubation, 10 µl of sperm-containing liquid was placed in the center of a Cell-Vu sperm counting cytometer. Sperm counts and motility were analyzed using the SCA®CASA system (Fertility Technology Resources) following the manufacturer’s instructions.

### 2.3. Preparation of single-nuclei suspensions and 10x Genomics libraries

Neonatal testes at PND6 were dissected and washed with HBSS. After removing the tunica albuginea, testicular parenchyma pooled from 3-4 pups from 3 litters per library were digested with 0.25% trypsin/EDTA containing 0.5 mg/ml DNase I for ∼15 min at 37 °C with gentle shaking and then quenched with 10% FBS. Suspensions were well pipette mixed and passed through a 70 μm and then a 40 μm cell strainer. Cell viability was evaluated and red blood cells were removed using a Red Blood Cell Lysis Buffer (BioLegend). THY1^+^ cells were isolated by magnetic labeling with anti-CD90.2 (THY1) MicroBeads (Miltenyi Biotec) following the manufacturer’s instructions. Nuclei isolation of THY1^+^ cells was performed according to the 10x Genomics protocol for single-cell Multiome analysis with modification. Briefly, THY1^+^ cell pellets (∼2 × 10^5^ cells) were incubated with 100 μl chilled 0.1× Lysis Buffer for 10 min on ice. Cell lysis was quenched by adding 1ml chilled wash buffer. Nuclei were harvested and ultimately resuspended with 1× nuclei buffer. The nuclei were counted after staining with trypan blue solution and then immediately processed for library preparation following the standard 10x Genomics Chromium Single Cell Multiome ATAC + Gene library preparation protocol.

### 2.4. Single nucleus (sn) Multiome data processing and analysis

#### 2.4.1 snMultiome data processing and quality control

In total 12 paired snRNA-seq and snATAC-seq libraries, including 2 replicates/group (6 samples) in the F1 generation and 1 replicate/group (3 samples) in each F2 and F3 generation, were sequenced on NovaSeq 6000 (Illumina). The FASTQ files were aligned to the UCSC mouse genome (mm10) and counted with cellranger-arc count (v2.0.0). A mean of 292,676 or, 233,904 reads per cell were sequenced for each snRNA or snATAC library, respectively (Supplementary Table S1). Datasets in each generation were aggregated by cellranger-arc aggr with depth normalization. RNA and ATAC matrices were imported to Seurat 4.4.0^68^ and Signac 1.11.9^69^, and data were separately analyzed by generation. Low-quality cells and multiplets were excluded using the following criteria: > 1000 genes and fragments, < 150000 UMI RNA and ATAC counts, TSS.enrichment > 1, nucleosome_signal > 1, %blacklist_fraction < 0.2 for F1 generation; > 1000 genes and fragments, < 75000 UMI RNA counts, < 200000 UMI ATAC counts, TSS.enrichment > 1, nucleosome_signal < 2, %blacklist_fraction < 0.2 for F2 generation; > 1000 genes and fragments, < 50000 UMI RNA counts, < 200000 UMI ATAC counts, TSS.enrichment > 1, nucleosome_signal < 2.5, %blacklist_fraction < 0.1 for F3 generation.

#### 2.4.2 snRNA-seq and snATAC-seq analysis

After quality control (QC), gene expression values from filtered snRNA-seq were log normalized, scaled, and dimensionally reduced by principal component analysis (PCA). Normalization of snATAC-seq data was performed with term-frequency inverse-document-frequency (TFIDF), followed by dimensional reduction via singular value decomposition (SVD) of the TFIDF matrix. After Harmony (v0.1.1) batch effect correction^70^, snRNA-seq and snATAC-seq data underwent uniform manifold approximation and projection (UMAP) analysis and subsequently constructed a weighted nearest neighbor (WNN) graph to leverage both modalities for the following visualization and clustering. Seurat clusters were annotated based on the expression of known cell-type markers from transcriptome profiles. Germ cells were subsetted, re-clustered, and imported into Monocle 3 (v1.3.1)^71^ for pseudotime trajectory analysis. Differentially expressed genes (DEGs) in germ cell subsets among treatment groups were determined by *FindMarkers* function (Wilcoxon rank sum test) in Seurat with *p.adjust* < 0.05, and used for Gene Ontology (GO) analysis with clusterProfiler (v4.2.2)^72,73^. The *CellCycleScoring* function of Seurat was used to compute cell cycle phases based on the expression of G2-M-and S-phase genes. For snATAC-seq, germ cell differentially accessible peaks (DAPs) were analyzed by *FindMarkers* function (LR test, *p.adjust* < 0.05). Detailed annotation of DAPs was performed with Homer (v4.11.1)^74^. For the epigenomic annotation of DAPs, publicly available ChIP-seq datasets of neonatal spermatogonia^59^ were used. The intersection of ChIP-seq peaks and DAPs was established utilizing Intervene^75^. DNA sequence motif information was obtained from the JASPAR database^76^, and BPA-or BPS-enriched motifs were identified using *FindMotifs* function in Signac. A per-cell motif activity score was further computed by running chromVAR^77^ with default parameters.

### 2.5. qRT-PCR

Total RNA was isolated from PND6 testes. The cDNA was synthesized with oligo (dT) primer from 1μg of total RNA using the High-Capacity cDNA Reverse Transcription Kit (Thermo Fisher). Relative gene expression was examined by Applied Biosystems™ SYBR Green Master Mix using a CFX Opus 96 Real-time PCR system (BioRad) as described previously^35,63^. Primer sequences were designed by NCBI’s design tools or from the PrimerBank database^78–80^ and are provided in Supplementary Table S11.

### 2.5. Immunohistochemistry

PND6 testes were fixed with 4% paraformaldehyde, paraffin-embedded, and sectioned (5 μm). Following previously established immunostaining of FOXO1 and STRA8^35,67^, the sections were deparaffinized, rehydrated, and treated by heat-induced antigen epitope retrieval in citrate buffer (pH 6.0) at 100 °C for 10 min. The sections were then blocked with 5% normal goat serum in TBS and immunolabeled with specific primary antibodies (Supplementary Table S12) overnight at 4 °C in a humidified chamber. The sections were washed with TBST and incubated with anti-rabbit biotinylated secondary antibody at room temperature for 30 min, followed by applying Avidin-Biotin Complex kits (Vector Laboratories) for 15 min at room temperature. The antigen signal was visualized by diaminobenzidine reaction and counterstained with hematoxylin. Images were obtained from a Leica DM4 B microscope, and FOXO1 and STRA8 positive cells were counted using Image J.

### 2.6. Statistical analysis

Statistical analyses for RT-qPCR and quantitative data for immunostaining were performed with GraphPad Prism (version 9.0.0). One-way ANOVA with post hoc Dunnett’s multiple comparisons test was used to determine the differences between control and BP-treated groups. A *P* value of <0.05 was considered statistically significant.

### 2.7. Data availability

The raw data of snMultiome sequencing has been deposited at NCBI/SRA (PRJNA1022459). We also created a cloudbased web tool (Webpage: https://kanakohayashilab.org/hayashi/bp/mouse/germcells/) for easy visualization of gene expression and ATAC peaks via the gene of interest searches.

## 3. Results

### 3.1. Prenatal exposure to BPA and BPS reduced sperm counts in the F1, F2, and F3 generations

We have previously reported that prenatal exposure to BPA or BPS reduced sperm counts and motility, and impaired staging of spermatogenesis in F1 and F3 males^35,63^. To confirm our prior results, we first examined sperm counts and motility in adult males across generations. As shown in Figure 1b, consistent with the previous findings, all BPA and BPS treatment groups exhibited significantly reduced sperm counts in the F1 to F3 generations on PND60 compared to the control group (CON). However, no significant differences were observed in sperm motility (Figure 1b), likely because the effects of BPA and BPS exposure were transmitted only paternally, unlike in our previous study where both maternal and paternal transmissions were involved^35^. Body weight, testis weight, and the testis-to-body weight ratio were also unaffected (Supplementary Figure 1a).

### 3.2. Single-cell transcriptomic and ATAC landscapes of spermatogonia in the F1 males

To further understand transgenerational defects on spermatogonial germ cells by ancestral BPA or BPS exposure, we performed snMulti-omics analysis (paired snRNA-seq and snATAC-seq from the identical nuclei) using neonatal PND6 spermatogonia. Across three generations, a total of 40,291 single nuclei were profiled. After QC filtering, 33,389 high-quality nuclei were retained, yielding averages of 3,666 genes and 7,965 peaks/nuclei (Supplementary Table S1). snRNA-seq and snATAC-seq datasets were paired by Seurat WNN analysis to construct a single UMAP embedding with both RNA and ATAC modalities for downstream analysis.

In the F1 generation, 16,591 nuclei were grouped into nine populations by the combined UMAP embedding, including Germ cells (*Ddx4*^+^ and *Dazl*^+^), Sertoli cells (*Sox9*^+^), Leydig cells (*Cyp17a1*^+^), Stromal cells (*Igf1*^+^ and *Pdgfra*^+^), Myoid cells (*Acta2*^+^), Macrophages (*Lyz2*^+^ and *Adgre1*^+^), Innate lymphocytes (*Il7r*^+^ and *Cd52*^+^), Pericytes cells (*Rgs5*^+^), and Endothelium cells (*Pecam1*^+^) (Figure 1c, Supplementary Figure 1b and Table S2). The cell type specificity was further validated with a peak-to-gene linkage analysis showing highly accessible chromatin states of known marker genes in each cell cluster based on snATAC-seq (Supplementary Figure 1c). A consistent UMAP clustering pattern was observed in the biological replicates of F1 samples (Supplementary Figure 1d), suggesting an unbiased capture of cell populations between treatments.

The germ cell population was further resolved to identify its subpopulations in different developmental states (Figure 1d, e, Supplementary Figure 2a, Supplementary Table S3), generating 4 clusters of SSCs (SSC1-4; *Id4^+^*, *Etv5^+^*, and *Gfra1^+^*), 1 cluster of progenitor-like cells (Progenitor; *Upp1*^+^, *Sox3*^+^, and *Rarg*^+^), and 3 clusters of differentiating spermatogonia (Diff1-3; *Kit^+^*, *Stra8^+^*, and *Sohlh1^+^*). Pseudotemporal trajectory analysis shows a clear developmental order from the cluster of SSC1 to Diff3 (Figure 1f and Supplementary Figure 2b), accompanied by the sequential expression of state-specific markers (Figure 1g). A similar trajectory pattern was observed between the control and BP treatment groups (Supplementary Figure 2c), indicating prenatal exposure to BPA or BPS did not disrupt the male germ cell developmental trajectory at the neonatal stage. Other newly identified cluster-specific marker genes and peaks were provided in Supplementary Tables S3 and S4. Therefore, snMulti-omics sequencing provides a high resolution of germ cell composition, enabling thorough exploration of the specific changes in germ cell development at the transcriptomic and epigenetic levels due to prenatal exposure to BPA and BPS.

### 3.3. Prenatal BPA or BPS exposure altered genes and biological processes associated with spermatogonial stem cell differentiation in the F1 generation

In the F1 spermatogonia, 6,842 up-regulated differentially expressed genes (up-DEGs) in the BPA group and 5,332 up-DEGs in the BPS group compared to the CON were identified (Figure 2a and Table S5). Notably, 4,388 (56.4%) up-DEGs overlapped between the two BP treatment groups (Figure 2a), with an overall higher expression in the BPA group (Figure 2b and Supplementary Figure 3a), suggesting BPA induced similar but stronger effects than BPS on neonatal germ cells. GO analysis showed that BPA and BPS exposure enhanced similar biological processes associated with epigenetic changes, energy metabolism, and apoptosis, as GO terms of “histone modification”, “methylation”, “ATP metabolic process”, “oxidative phosphorylation”, and “intrinsic apoptotic signaling pathway” were enriched (Figure 2c). In addition, mitotic and meiotic cell cycle processes were also enhanced. Further trajectory analysis showed that most of the genes involved in these enriched pathways gradually up-regulates their expression along the differentiation path (Figure 2d). Core genes for pathways of oxidative phosphorylation (OXPHOS), apoptosis, mitosis, or spermatogonial differentiation/meiosis were confirmed by qRT-PCR in independent samples of the testis, and most of these genes were consistently enhanced or showed a tendency (*P* < 0.1) to increase in the BPA and/or BPS groups (Figure 2e). Moreover, compared to BPS, BPA exposure enhanced biological processes of “oxidative phosphorylation”, “ATP metabolic process”, “regulation of translation”, and “cell cycle phase transition”, indicating that BPA exposure induced more changes in metabolism and cell cycle progression than BPS (Supplementary Figure 3).

**Figure 2.**
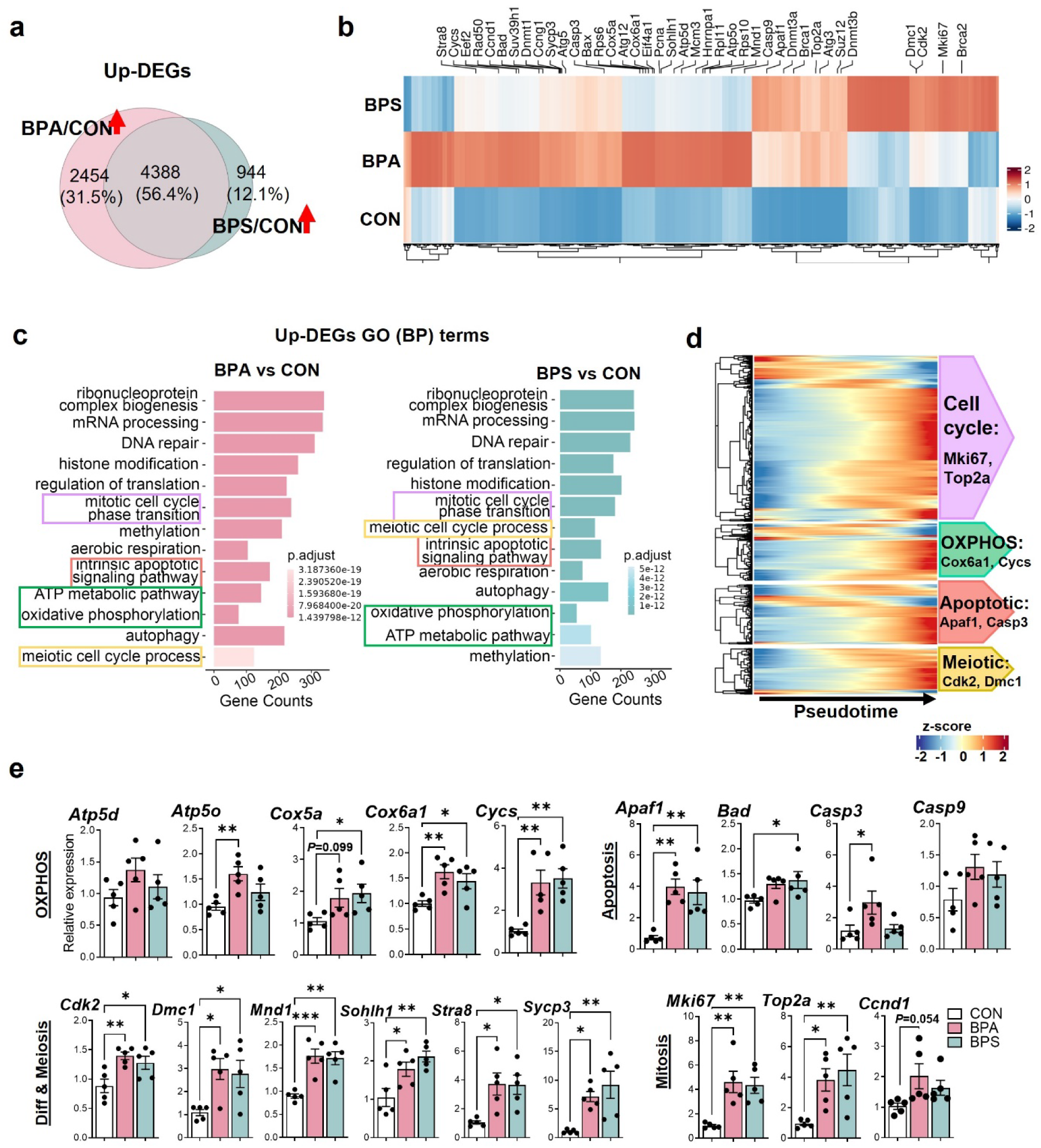
Genes and biological processes up-regulated by prenatal BPA and BPS exposure in the F1 germ cells. (a) Venn diagram shows the numbers and the overlaps of genes up-regulated by exposure to BPA (BPA/CON) and BPS (BPS/CON). (b) Heatmap shows the up-regulated genes in each treatment group. (c) Enriched GO biological processes in germ cells by prenatal BPA and BPS exposure. (d) Heatmap shows the expression pattern of genes for enriched pathways alongside the pseudo-developmental process of germ cells. (e) Verification of differential gene expression by RT-qPCR. \**P* < 0.05, \*\**P* < 0.01, \*\*\**P* < 0.001, mean ± SEM, n = 5/group.

As for the down-regulated DEGs (down-DEGs), 433 and 172 genes were identified in the BPA and BPS groups, respectively, and 114 genes of them overlapped, including *Mcam*, *Ret*, *Cd9*, *Egr1*, *Hmgcr*, and *Pde1c* (Figure 3a, 3b, and Supplementary Table S5, *p.adjust* <0.05). GO analysis suggests that both BPA and BPS exposure weaken biological processes associated with cellular responses to internal and external stimuli, including response to amino acid, acid chemical, calcium ion, mechanical stimulus, and wounding. GO terms associated with response to glucose/hexose/hormone and TGFβ signaling were uniquely enriched in the BPA group, whereas biological processes related to macrophage activation and cytokine production are enriched in the BPS group (Figure 3c). Notably, most down-DEGs showed a higher expression in the early stage of germ cell development trajectory and decreased gradually after that (Figure 3d), suggesting their potential functions for the maintenance of stemness. Representative genes were validated using qRT-PCR in independent samples, and their expression patterns mostly agreed with the sequencing results (Figure 3e).

**Figure 3.**
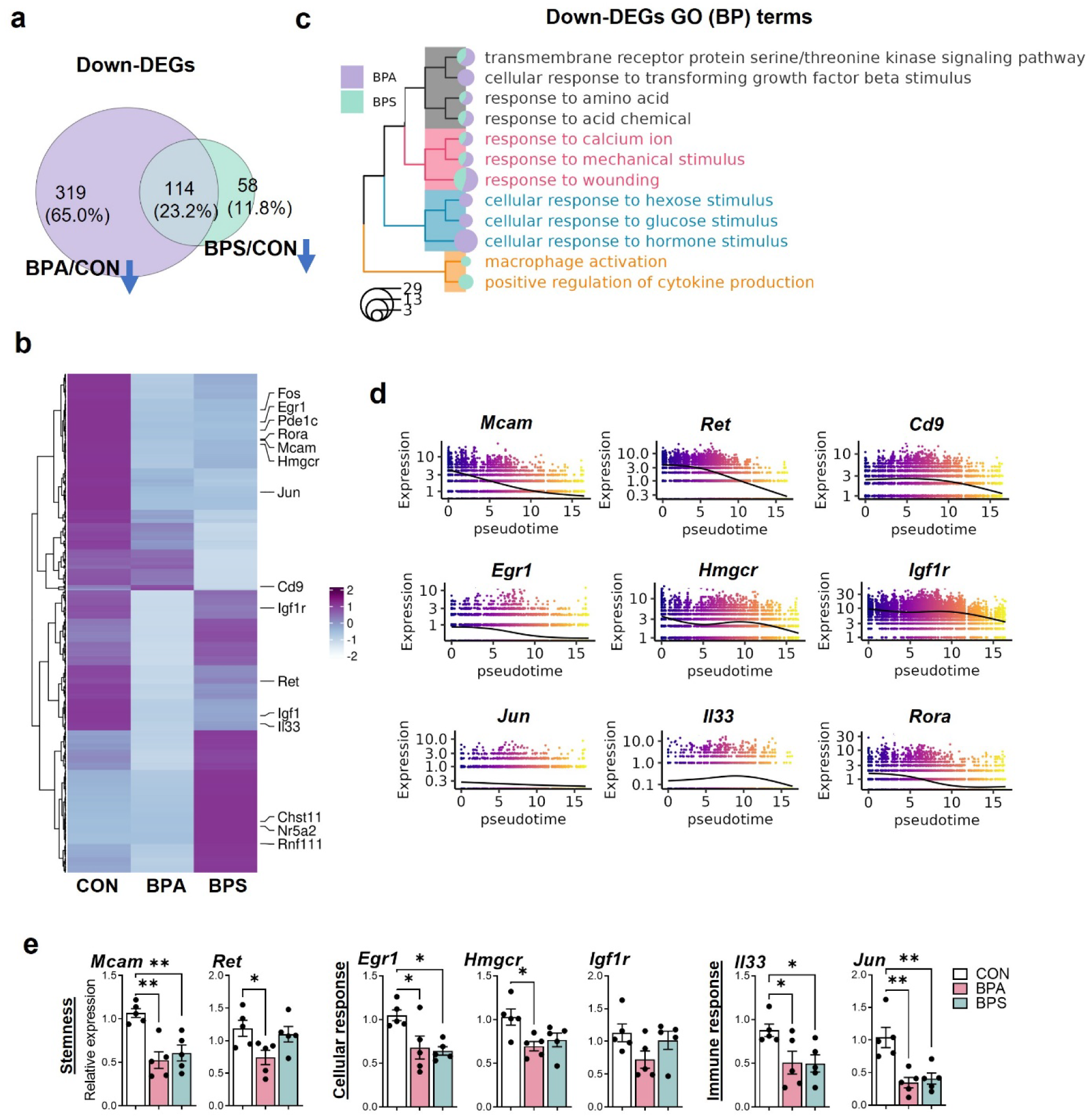
Down-regulated genes and biological processes by prenatal BPA and BPS exposure in the F1 germ cells. (a) Venn diagram shows the numbers and the overlaps of genes down-regulated by exposure to BPA (BPA/CON) and BPS (BPS/CON). (b) Heatmap shows the down-regulated genes in each treatment group. (c) Treeplot shows the hierarchical clustering of enriched biological processes. (d) Expression pattern of representative down-regulated genes alongside the pseudo-developmental process of germ cells. (e) Verification of differential gene expression by RT-qPCR. \**P* < 0.05, \*\**P* < 0.01, mean ± SEM, n = 5/group.

In summary, transcriptomic analysis of germ cells suggested that prenatal exposure to both BPA and BPS promoted gene expression for germ cell differentiation while reducing gene expression for stemness maintenance, which might disrupt the spermatogonial homeostasis in the F1 generation.

### 3.4. Prenatal exposure to BPA and BPS imbalanced spermatogonial stem cell differentiation in the F1 testis

Cell cycle scoring was conducted to show the differences in the cell cycle progression of germ cells between treatments in the F1 generation. While the distribution pattern of cell cycle phases was consistent between all three groups, more cells in phases of synthesis (S) and gap 2 (G2)-mitosis (M) were found in the BPA and BPS treatment groups, suggesting enhanced potential of differentiation (Figure 4a and 4b). Furthermore, the numbers of germ cell subpopulations were quantified to calculate their relative proportions (Figures 4c and 4d). Consistently, more proportions of progenitor and differentiating cells and fewer SSCs were observed in the BP treatment groups compared to the CON (Figure 4d). To verify the effects of BPA and BPS exposure on germ cell differentiation, we examined undifferentiated and differentiating cell proportions in F1 neonatal testis. On PND6, FOXO1 and STRA8 were immunostained, as markers for undifferentiated and differentiating cells, respectively. Significantly reduced % of FOXO1^+^ tubules and increased STRA8^+^ cells per positive tubule were observed following BPA and BPS exposure (Figure 4e and 4f), whereas FOXO1^+^ cells per positive tubule and % of STRA8^+^ tubules were comparable between control and BP exposure groups. Consequently, in line with our transcriptomic findings, prenatal BPA and BPS exposure elevated the proportion of differentiating spermatogonia, during which the epigenetic alterations likely program these long-lasting effects.

**Figure 4.**
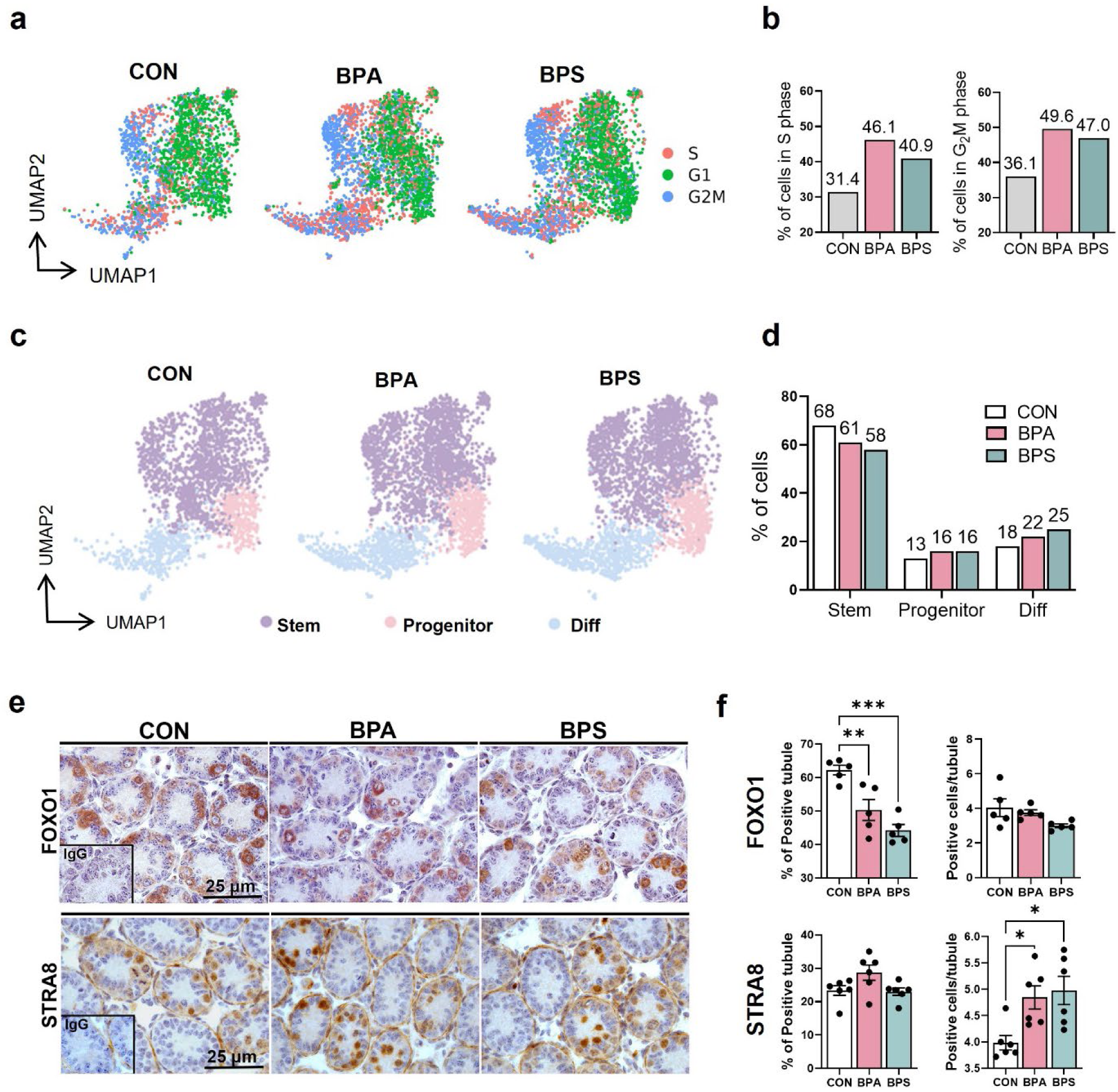
Cell cycle progression and differentiating states of germ cells in F1 germ cells. (a) UMAP distribution of cell cycle phases. (b) The proportions of germ cells in S and G_2_M phases. (c) UMAP visualization of the distribution of SSCs, progenitors, and differentiating cells. (d) Bar plot shows the cell proportions in three stages of germ cell differentiation. (e) Immunohistochemistry analysis of PND6 testis sections stained with FOXO1 or STRA8. (f) The percentages of FOXO1^+^ and STRA8^+^ tubules and positive cells per tubule. \**P* < 0.05, \*\**P* < 0.01, \*\*\**P* < 0.001, mean ± SEM, n = 5/group. SSC, spermatogonial stem cell.

### 3.5. Prenatal exposure to BPA and BPS changes chromatin accessibility and TF motif activity associated with germ cell differentiation in the F1 generation

The chromatin accessibility of germs cells in the F1 generation was further analyzed to understand the epigenetic changes caused by prenatal BPA and BPS exposure. Of 170,123 total ATAC peaks, 4,729 and 2,931 differential accessible regions (DAPs) were identified in the groups of BPA and BPS, respectively, compared to the CON group (Figure 5a and Table S6). Among them, 1,697 DAPs of the BPA group correspond to 1,591 up-DEGs and 106 down-DEGs at the transcriptomic level (Figure 5a). For BPS exposure, 914 DAPs match with 886 up-DEGs and 28 down-DEGs. This finding suggests that the opening status of chromatin does not always positively correlate with gene expression. In addition, these DAPs were predominately located in promoter regions (Figure 5b), suggesting potential programming of transcriptional activity changes caused by BP exposure. Interestingly, consistent with our transcriptomic results, which suggest stronger disturbance caused by exposure to BPA than BPS, 47.45% of DAPs are located in the promoter region of germ cells with gestational BPA exposure, but this number reduced to 38.96% in the BPS group (Figure 5b).

**Figure 5.**
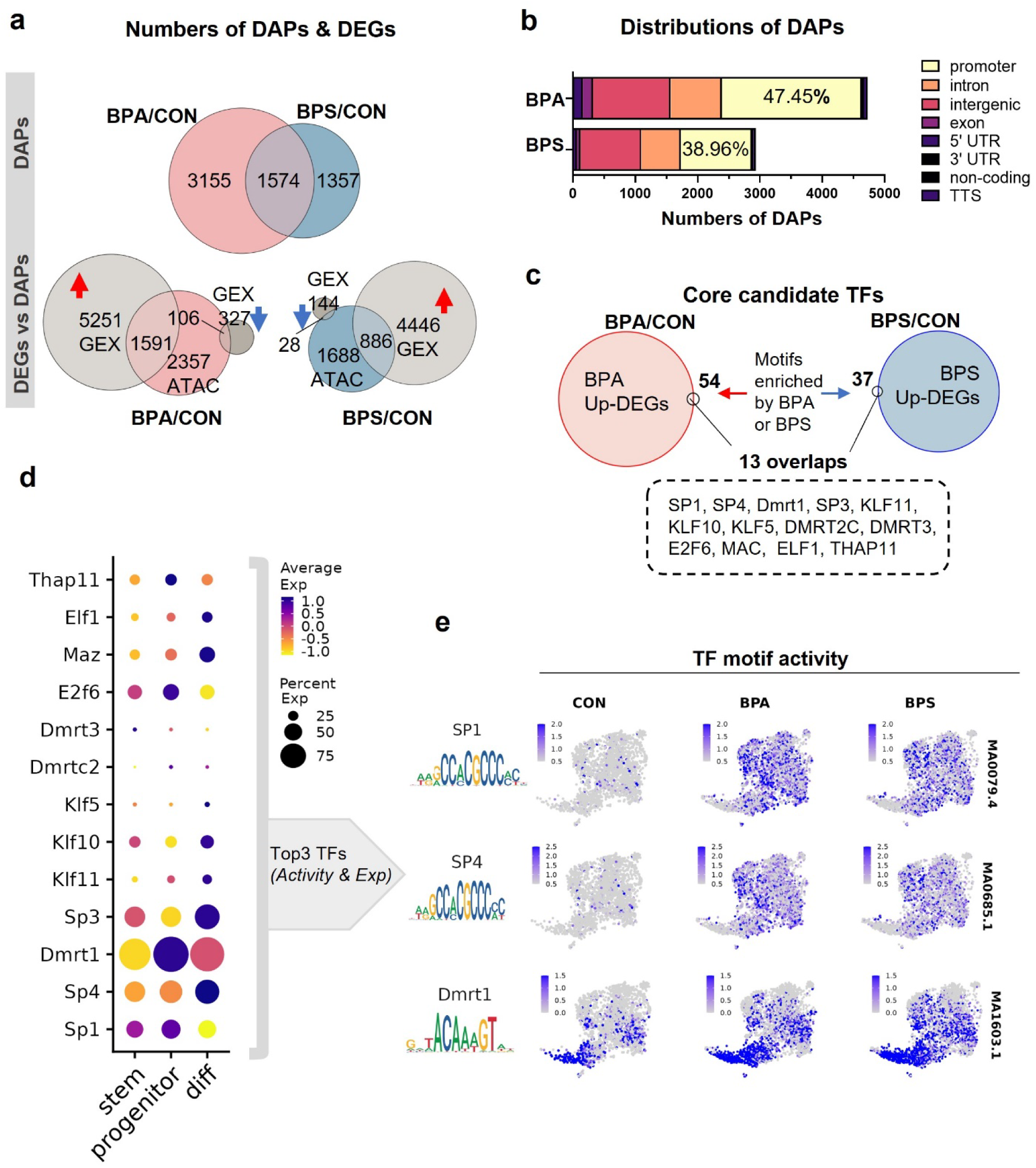
Effects of prenatal exposure to BPA and BPS on chromatin accessibility of germ cells in the F1 generation. (a) Top panel, differentially accessible peaks (DAPs). Bottom panel, the overlapping genes between DAPs (ATAC data) and DEGs (GEX data). (b) The genomic distribution of DAPs caused by gestational BPA and BPS exposure. (c) Core candidate TFs were acquired by intersecting the DEGs with enriched TF motifs. (d) Dot plot shows the gene expression pattern of acquired core TFs in different stages of germ cell differentiation. (e) Motif sequences (left) and UMAP visualization of TF chromVAR deviations (right) of SP1, SP4, and DMRT1 between groups. TF, transcriptional factor.

The motif enrichment analysis was conducted using the Signac package to obtain the lists of enriched TF motifs. By intersecting them with the up-regulated genes in each BP treatment group, 13 overlapped potential core TFs were identified in the BPA and BPS groups (Figure 5c). It was noted that most of these TFs exhibit stronger expression in the Progenitor and Diff populations than in the SSCs (Figure 5d). Then, the motif activity of these TFs was computed by chromVAR, and the results showed that the activity of SP1, SP4, and DMRT1, was all enhanced in the BPA and BPS groups (Figure 5e). To better understand the long-lasting effects caused by gestational BPA and BPS exposure, the downstream signaling transduction pathways of the candidate TFs of SP1, SP4, and DMRT1, need to be explored.

### 3.6. Signal transduction of SP1, SP4, and DMRT1 in germ cells drives the differentiation process

To reveal the possible target genes of the candidate TFs, we established a prediction framework (Figure 6a). First, we overlapped the entire mouse promoter regions with total ATAC peaks and detected 18,410 peaks in the promoter regions (named “promoter peaks”). Then, we scanned which genes annotated to the promoter peaks that contain the TF motif sequences of interest. Since SP1 and SP4 belong to the SP family and share a similar motif sequence, we examined them together and only included genes with both motifs for the downstream analysis. Lastly, with the advantage of multi-omics sequencing, the gene lists were further narrowed by intersecting the genes containing interested motifs with DEGs from snRNA-seq data.

**Figure 6.**
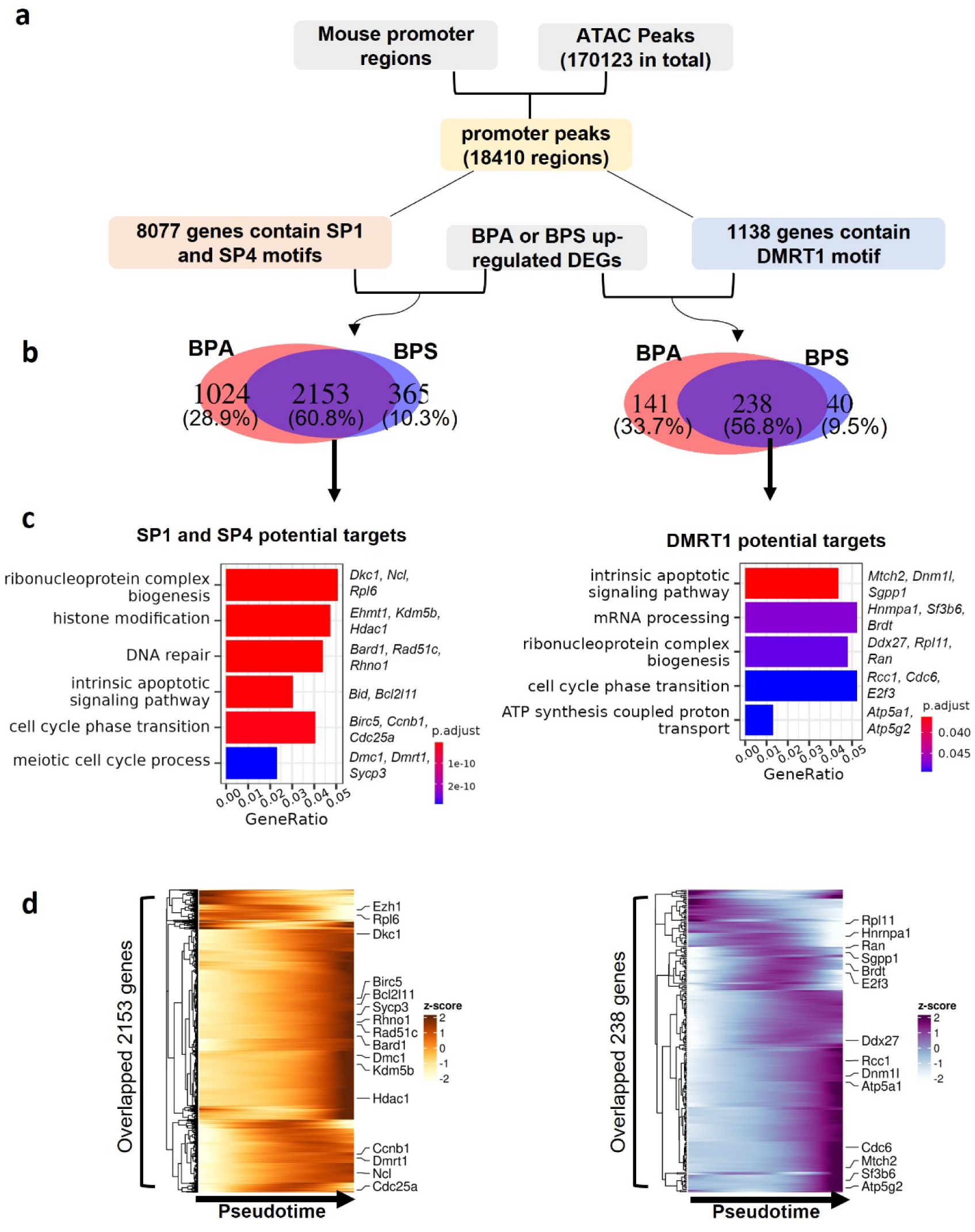
Identification of potential target genes of SP1/SP4 and DMRT1 in the F1 generation. (a) Workflow of the analysis framework. (b) Venn diagrams reveal the numbers and the overlaps of up-regulated genes potentially targeted by SP1/SP4 or DMRT1. (c) GO enrichment analysis of predicated target genes. (d) Heatmap shows the expression pattern of target genes along the pseudotime trajectory.

As shown in the Venn diagrams, 3,556 up-DEGs were potentially regulated by SP1/SP4 and DMRT1 in the BPA group, and the number is 2,796 in the BPS group (Figure 6b and Supplementary Table S9). Within either SP1/SP4 or DMRT1 potential targets, ∼60% of overlapping was observed between BPA and BPS groups (Figure 6b). GO analysis of these overlapping genes showed enrichment in the processes of ribonucleoprotein complex biogenesis, histone modification, intrinsic apoptosis, DNA repair, cell cycle phase transition, or ATP synthesis (Figure 6c). In addition, a pseudotime-ordered, gene expression heatmap was used to investigate their expression patterns along the differentiation trajectory of germ cells. Expression of these genes was mostly enhanced in the middle and late stages of the differentiation process (Figure 6d).

Using the same strategy, genes potentially down-regulated by SP1/SP4 and DMRT1 were also analyzed, and the number was significantly fewer compared to the up-regulated genes (Supplementary Figure 4a). A total of 190 down-DEGs (175 BPA and 52 BPS) were predicated to be downstream targets of SP1/SP4 (Supplementary Figure 4b). Only 37 genes, including those correlated to stemness (e.g. *Mcam*, *Ret*) and cfos/cJun components (e.g. *Fosb* and *Junb*) overlapped in BPA and BPS groups. For DMRT1, 28 and 6 down-regulated genes were filtered out in the BPA and BPS groups, respectively, with 5 overlaps of *Aff3*, *Fosb*, *Glis3*, *Ltbp4*, and *Mov10l1* (Supplementary Figure 4b). These results suggested that SP1, SP4, and DMRT1 were involved in enhancing the processes associated with neonatal germ cell differentiation via regulating multiple downstream gene sets with consistent functions for differentiation programming.

### 3.7. Comparation of transcriptomic changes of germ cells with prenatal exposure to BPA and BPS across F1 to F3 generations

To understand the transgenerational effects of BPA and BPS exposure in spermatogonia, snMulti-omic results from the F2 and F3 generations were compared. Similar to the F1 generation, we observed 9 major cell types, including a majority of germ cells (∼50%) and several somatic cells from 5,334 or 11,410 high-qualified nuclei in the F2 and F3 THY1^+^ testicular cells, respectively (Supplementary Figure 5a and Table S2). An unbiased capture of cell populations was confirmed among biological replicates in each generation (Supplementary Figure 5b). Three main subtypes of germ cells, including SSCs, progenitors, and differentiating cells, were classified in the F2 and F3 generations as those of F1 (Figure 7a, Supplementary Figure 5c, and Table S3). An additional SSC cluster “SSC5” was identified in the BPA and BPS groups of the F2 and F3 generations (Supplementary Figure 6a). GO analysis showed that genes enriched in SSC5 are related to biological processes of p53-mediated signal transduction and/or DNA damage response in addition to pathways of “cytoplasmic translation”, “ribosome biogenesis”, and “cell cycle phase transition” (Supplementary Figure 6b and 6c). Unlike F1, the proportions of SSCs, progenitor, and differentiating cells were comparable between groups of treatments (Supplementary Figure 6d).

**Figure 7.**
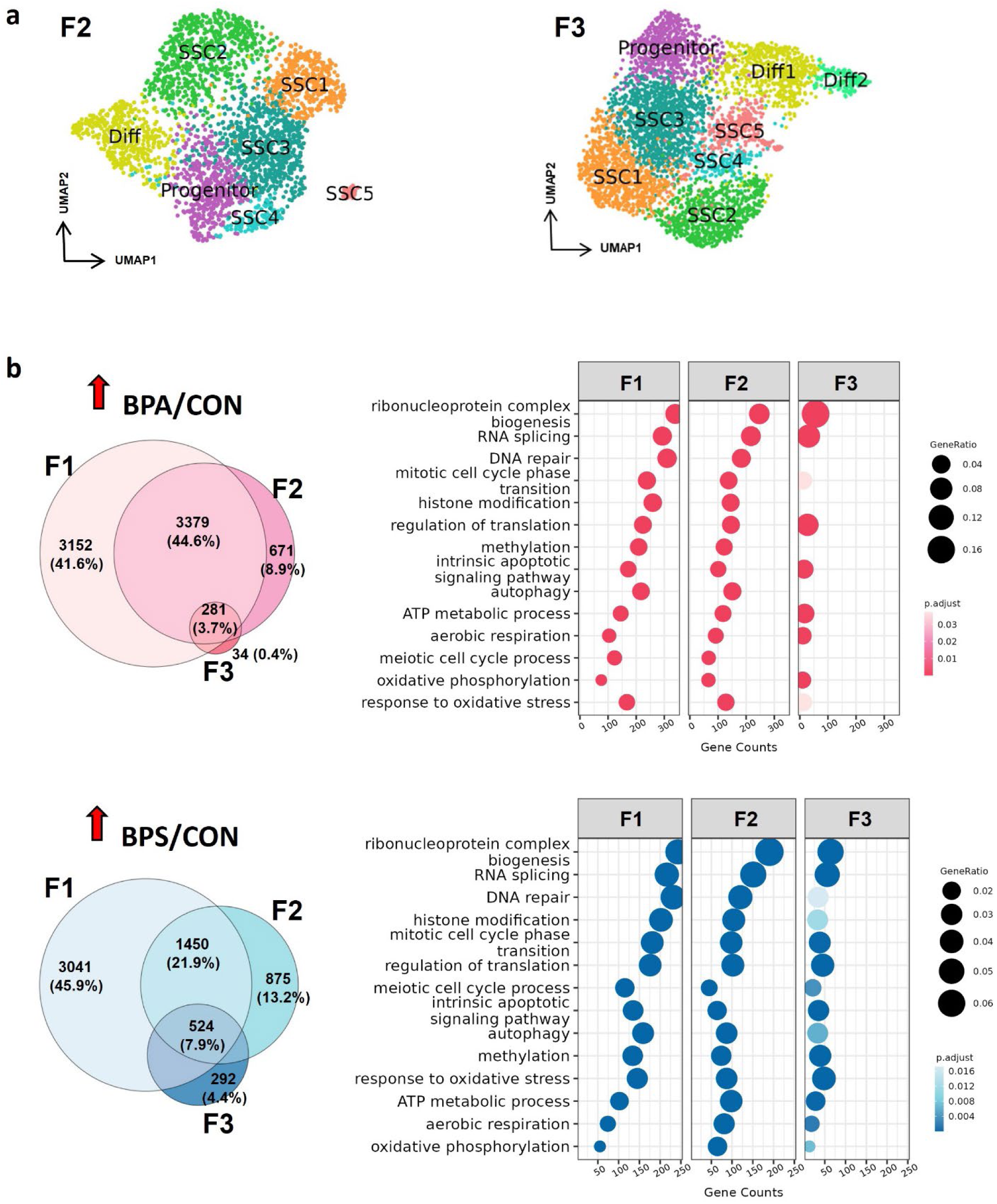
Transcriptomic changes on neonatal germ cells across generations caused by prenatal exposure to BPA and BPS. (a) UMAP plots of germ cell sub-clusters in the F2 and F3 generations. (b) Up-regulated genes and enriched GO terms in F1, F2, and F3 germ cells exposed to BPA or BPS.

Next, we analyzed the up-regulated genes in BP treatment groups compared to the CON across F1, F2, and F3 generations. Strikingly, in either the BPA or BPS group, the DEGs highly overlapped between the F1 and F2 generations, but the numbers greatly decreased in the F3 generation (Figure 7b). As a result, only small numbers of genes were consistently up-regulated throughout all three generations, including 281 genes in the BPA group, and 524 genes in the BPS group (Figure 7b). In agreement with DEGs results, GO terms of “DNA repair”, “histone modification”, “methylation”, “autophagy”, and “meiotic cell cycle process” were not enriched in the F3 generation of the BPA exposure group. In contrast, biological processes up-regulated by BPS exposure in the F1 germ cells were consistently enhanced in both F2 and F3 generations (Figure 7b). Therefore, compared to BPA, the effects of prenatal exposure to BPS on transcriptomes of germ cells appeared to be more transgenerationally sustained.

As for the downregulated genes caused by F0 BPA exposure, 433, 129, and 2399 genes were identified in the F1, F2, and F3 generations, respectively (Supplementary Figure 7a). It is noted that the number of BPA down-regulated genes increased greatly in the F3 generation. Interestingly, GO terms enriched by these F3 down-DEGs included the discontinued up-regulated processes in the F3 generation (Supplementary Figure 7a). In the groups of BPS, we observed comparable numbers in down-DEGs between generations. However, the gene sets and their enriched GO terms were distinct among the F1, F2, and F3 generations (Supplementary Figure 7b).

### 3.8. Transgenerational impacts of chromatin accessibility landscapes in germ cells of the F1, F2, and F3 generations

BPA and BPS exposure of the F0 females resulted in comparable numbers of DAPs with similar genomic distribution patterns in germ cells of the F1 and F2 generations (Figure 8a and Supplementary Table S7). However, this number largely decreased in the F3 generation along with dramatically reduced distribution in the promoter regions (Figure 8a and Supplementary Table S8). In addition, annotation of DAPs using publicly available ChIP-seq datasets for PND6 spermatogonia^59^ revealed that approximately 80% of BPA and BPS DAPs in the F1 and F2 generations overlapped with histone post-translational modifications that are important for transcriptional activation including H3K4me1/2/3 and H3K27ac, whereas only a few (∼20%) overlapped with histone repressive modifications such as H3K9me2/3 and H3K27me3. However, in the F3 generation, BPA and BPS DAPs showed much lower intersection (∼50%) with active histone marks, but higher (∼30%) overlapping with repressive marks especially H3K9me3 than those in the F1 and F2 generations. Therefore, our data suggests differential chromatin accessibility changes caused by direct exposure of germ cells to BPA and BPS for the F1 and F2 generations and indirect exposure for the F3 generation.

**Figure 8.**
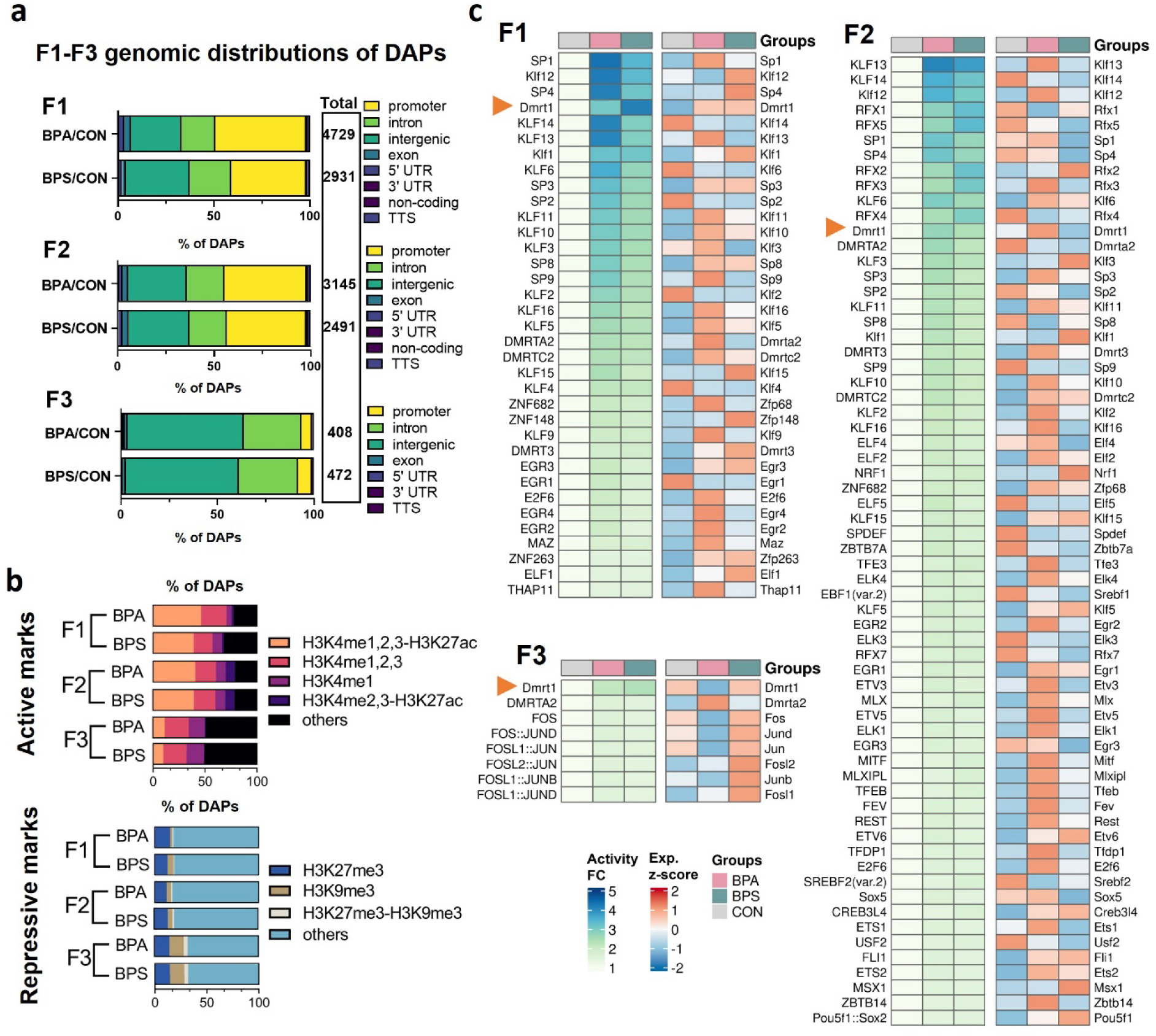
Changes in chromatin accessibility and histone modifications of neonatal germ cells across 3 generations with F0 prenatal exposure to BPA and BPS. (a) The genomic distribution of DAPs in germ cells of the F1, F2, and F3 generations. (b) Bar plots showing the overlaps of identified DAPs with genomic regions significantly enriched for repressive or active histone marks. (c) Heatmaps showing the activity of TF motifs and their associated gene expression levels between groups in the F1, F2, and F3 generations. FC, fold change. Exp, gene expression.

Using a combined motif enrichment (Signac) and activity analysis (ChromVAR), we identified lists of TFs that might be activated by BPA or BPS exposure throughout all three generations (Supplementary Table S10). The fold change (FC) of TF activities and their associated encoded gene expression levels were displayed by heatmap (Figure 8c). Among them, the TF activities and gene expression levels of DMRT1 were consistently enhanced throughout F1 to F3 generations in both the BPA and BPS groups (Figure 8c). Considering the downstream targets of DMRT1 illustrated in Fig. 6 and their corresponding functions for germ cell programming, changes in DMRT1 motif activities and gene expression levels might be a key factor for the disturbance of germ cell development, which may account for reduced sperm counts in 3 consequent generations with F0 prenatal BPA or BPS exposure.

## 4. Discussion

EDCs like BPA and phthalates are pervasive in our environment and have sparked serious concerns about their potential impacts on human health^4,81,82^. Studies on animals suggest that exposure to EDCs during development can not only affect the exposed individuals but also have repercussions on their offspring^11,17,58,83^, making EDC contamination a serious environmental issue. It is currently believed that the genetic changes caused by environmental factors are passed down through the germline, however, the molecular mechanisms of transgenerational inheritance are not fully elucidated^5,51,84^. In this study, we employed snMulti-omics to investigate how prenatal exposure to BPA and BPS affects the transcriptome and chromatin accessibility in the germline of male mice across three generations. Our work confirmed that prenatal exposure to an environmentally relevant low-dose BP negatively affects sperm counts across three generations and provided novel insights into the dynamic changes of the transcriptome and chromatin accessibility landscapes in germ cells.

In males, the development of early spermatogonia follows a well-defined and unique trajectory that is critical for maintaining the SSC pool and ensuring successful spermatogenesis^47,85^. Several in vivo and in vitro studies reported the cytotoxic effect of BPA exposure on SSCs, leading to compromised survival of SSCs and increased apoptosis^86–91^. Our results showed that prenatal exposure to BPA and BPS at a dose of 50 µg/kg/b.w./day led to an increased proportion of SSCs undergoing differentiation but a reduced proportion of undifferentiated cells within the F1 germ cell population, suggesting a disrupted balance between the stemness and differentiation of SSCs. Consistently, our transcriptional analysis revealed that BP exposure up-regulated genes and biological processes associated with spermatogonial differentiation, such as genes related to meiosis regulation (e.g., *Stra8*, *Sohlh1/2*, and *Sycp3*), oxidative phosphorylation, and cell cycle processes. However, the imbalance between undifferentiation and differentiation was not obvious in the F2 and F3 generations.

Epigenetic changes such as DNA methylation and histone modifications in the testis were associated with impaired reproductive capacity^92^. However, epigenetic changes of neonatal spermatogonia exposed to BPs have not been well documented. The impacts of prenatal exposure to BPA and BPS on chromatin states of germ cells across three generations were analyzed at the single-cell level in this study. BP exposure was found to mainly affect the TF activities of spermatogonia, as the majority of DAPs were located in the promoter regions. Furthermore, we identified 13 transcription factors potentially affected by BP exposure, including members of the SP/KLF family. SP1 and SP3 are known to regulate the gene expression of DNA methylation-related enzymes, including *Dnmt1*, *Dnmt3a*, and *Dnmt3b*^93,94^.

Interestingly, DMRT1 motif activity was consistently elevated in both BPA and BPS-exposed groups throughout all three generations. DMRT1 plays multiple pivotal roles in perinatal germ cell development by governing sex determination, maintaining the germ cell lineage, and ensuring proper differentiation^95–97^. Aberrant activation of DMRT1 leads to dysregulated gene expression during these critical developmental processes, which are likely responsible for the disrupted spermatogonial activities across generations observed in this study. Similar changes at both transcriptomic and ATAC levels caused by F0 BPA and BPS exposure were found between the F1 and F2 generations but not with the F3 generation. These results suggest that the epigenetic alterations acquired from ancestor exposure to BPA and BPS were not consistently inherited between generations. In line with this finding, distinct patterns of differentially methylated regions (DMRs) in sperm have been reported between generations following ancestral exposure to EDCs^98,99^. Future research uncovering the mystery of epigenetic regulatory mechanisms for germ cell development is sorely needed to better understand the transgenerational effects caused by ancestor BP exposure.

In summary, our work found that the environment-relevant dose of BPA and BPS exposure during gestation induces dramatic epigenetic changes in germ cells, disrupting their balance between undifferentiation and differentiation, during which the transcription factor DMRT1 might play a key role. While further research is necessary to fully understand the signaling transduction mechanism of epigenetic changes-induced long-term effects on reproductive defects in offspring, our study offers detailed information about changes in chromatin accessibility alongside the gene expression profiles of individual germ cells throughout multiple genes.

## Author contributions

M.S. and K.H. designed the research; L.Z and M.S. performed research, analyzed data, and wrote the paper; K.H. reviewed and revised the paper; S.W. configured the computer system and established the environment necessary for data analysis; J.A.M. provided critical feedback on the manuscript; all authors read, reviewed, and approved the manuscript.

## Acknowledgment

This work was supported by NIH/NIEHS R21 ES031607. We thank Dr. Nathan C Law for help with coding issues. We also thank Madeleine Harvey, Logan Butler, and Esther Langholt for their help with tissue collection.

