## Supplemental Figures for "Environmentally-relevant doses of bisphenol A and S exposure in utero disrupt germ cell programming across generations resolved by single nucleus multi-omics"

**Supplementary Figures:**

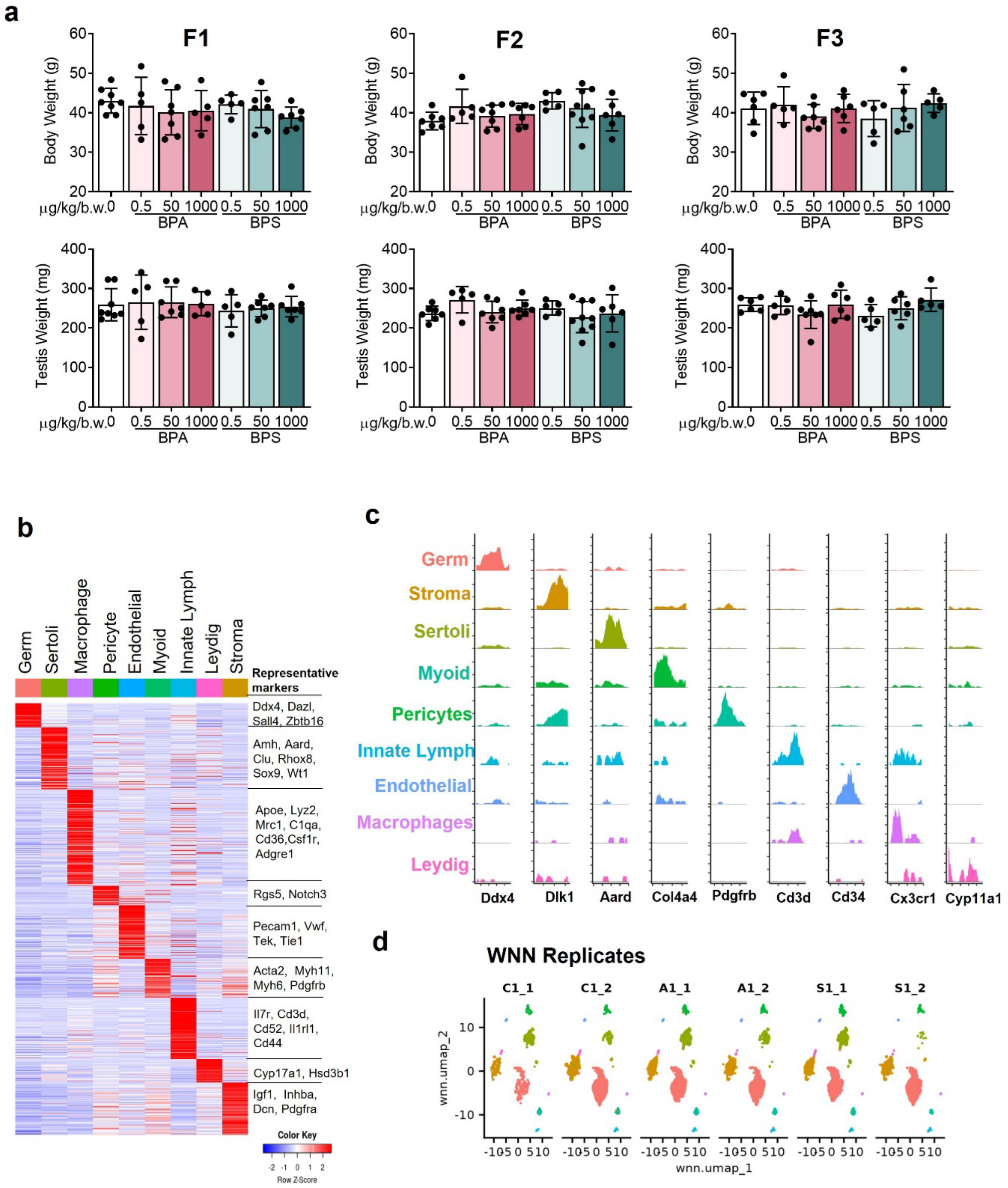


**Supplementary Figure 1**. Phenotypic characterization and identification of cell-specific markers in the F1 testis. (a) Body and testis weights. (b) Heatmap showing identified representative marker genes for each cluster. (c) Coverage plot shows the ATAC peaks of specific markers for each cell type. (d) UMAP visualization of cell distribution between biological replicates.

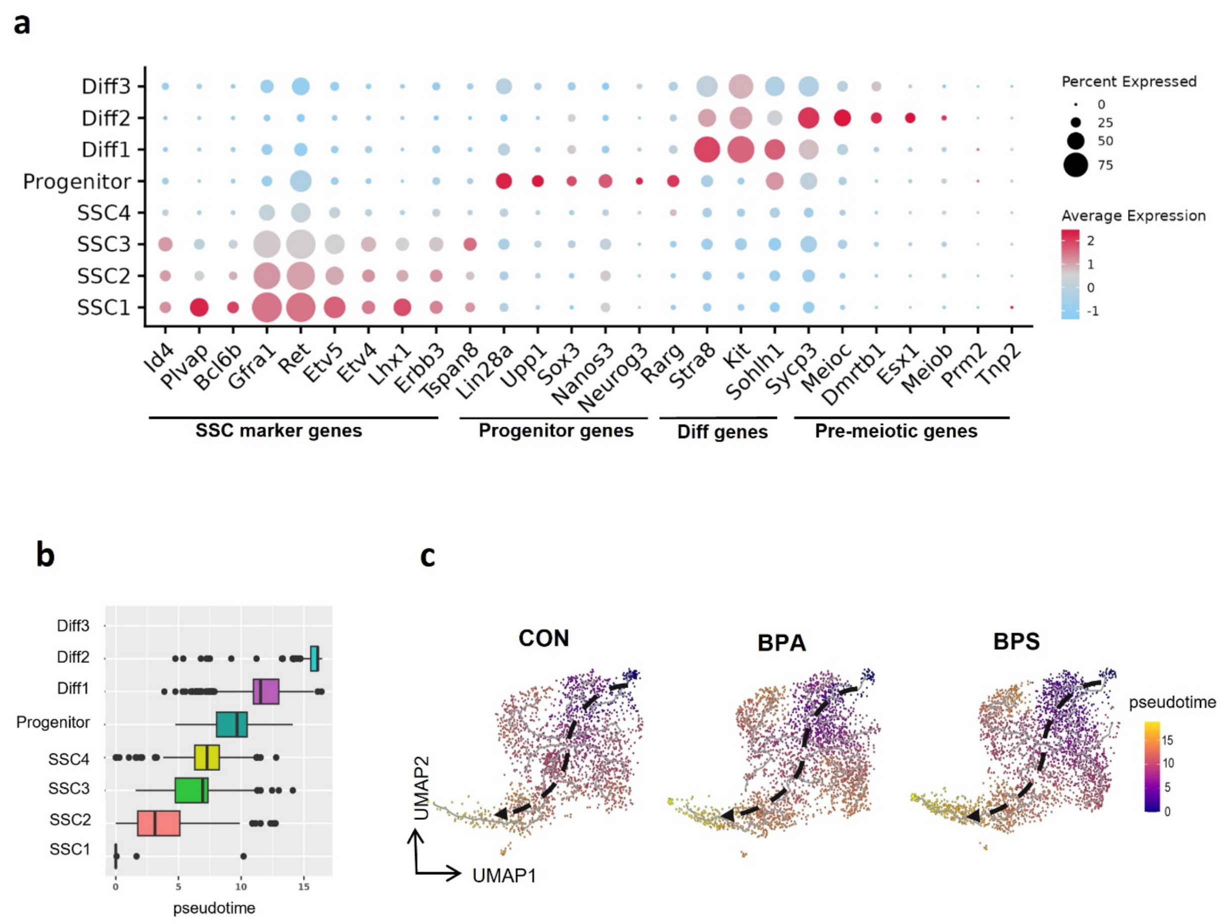


**Supplementary Figure 2**. Transcriptomic analysis in the F1 germ cells. (a) Dot plot depicts the scaled average expression of selected marker genes within each cluster in the F1 germ cells. (b) Box plot indicates the order of pseudotime trajectory by clusters. (c) Comparison of monocle pseudotime trajectory analysis between groups of different treatments.

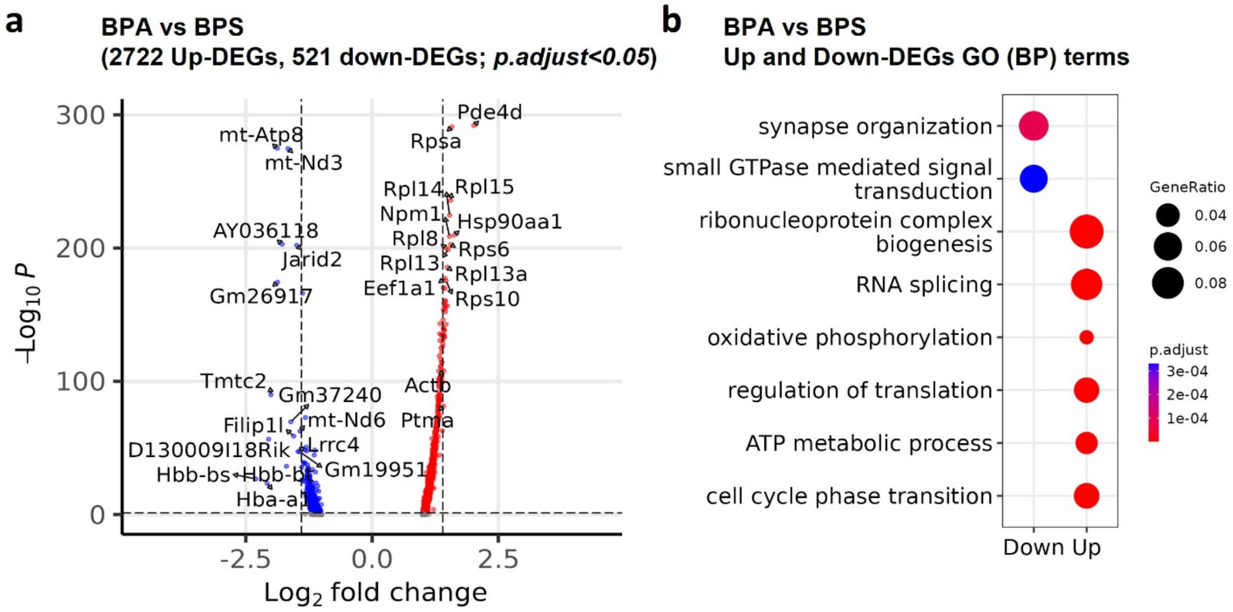


**Supplementary Figure 3**. Characterization of differentially expressed genes (DEGs) between BPA and BPS exposure groups in the F1 generation. (a) Volcano plot of up-regulated and down-regulated DEGs: BPA vs BPS. (b) Dotplot depicts the GO annotation of down- and up-regulated DEGs.

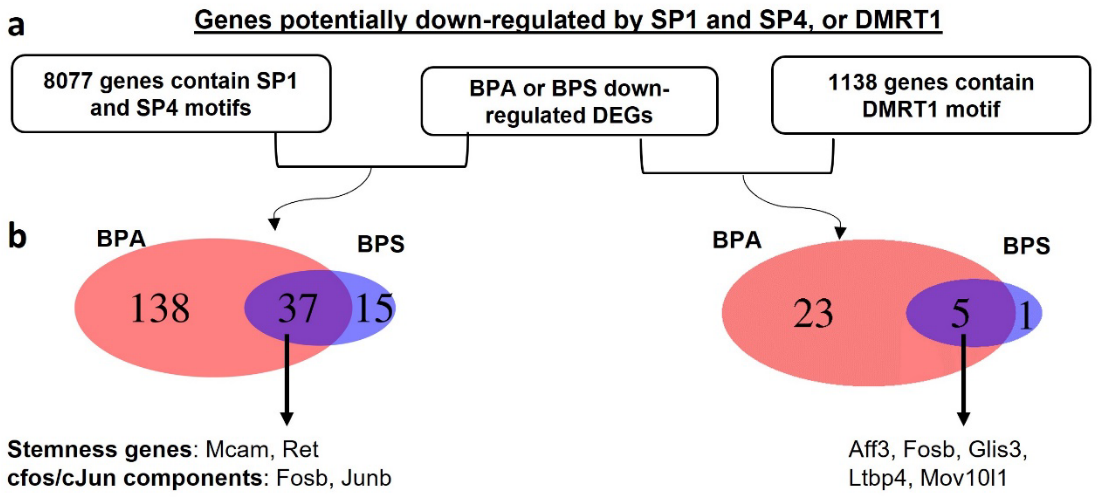


**Supplementary Figure 4**. Identification of genes potentially down-regulated by SP1/SP4 and DMRT1 in the F1 generation. (a) The framework of identification. (b) Venn diagrams show the numbers and the overlaps of down-regulated genes potentially targeted by SP1/SP4 and DMRT1.

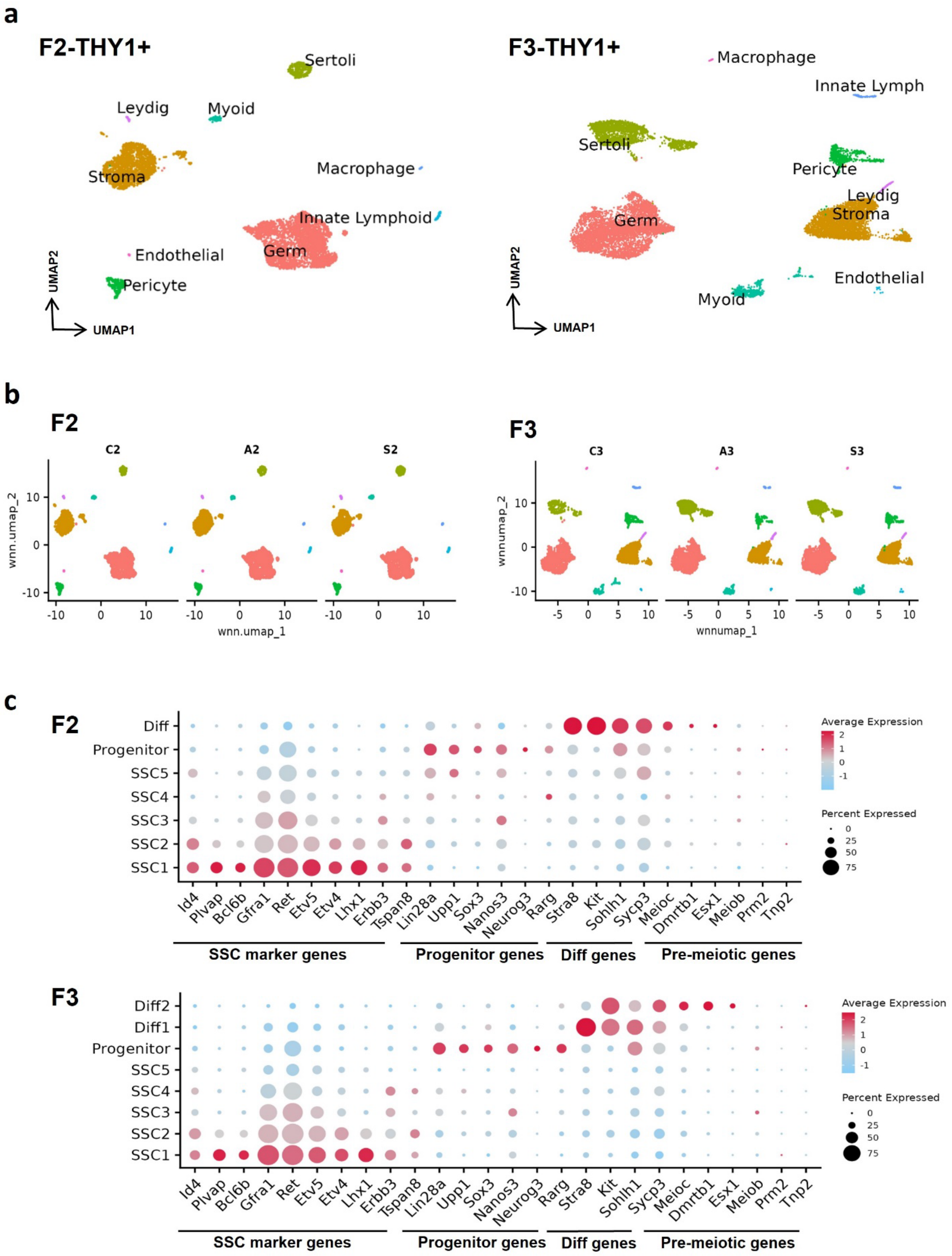

a
F2-THY1+
Sertoli
Leydig
Myoid
Stroma
Macrophage
Innate Lymphoid
Endothelial
Germ
Pericyte
UMAP2
UMAP1
F3-THY1+
Macrophage
Innate Lymph
Sertoli
Pericyte
Leydig
Stroma
Germ
Endothelial
Myoid
UMAP2
UMAP1
b
F2
C2
A2
S2
wnn.umap_2
wnn.umap_1
F3
C3
A3
S3
wnnumap_2
wnnumap_1
c
F2
Diff
Progenitor
SSC5
SSC4
SSC3
SSC2
SSC1
Id4 Plvap Bcl6b Gfra1 Ret Etv5 Etv4 Lhx1 Erbb3 Tspan8 Lin28a Upp1 Sox3 Nanos3 Neurog3 Rarg Stra8 Kit Sohlh1 Sycp3 Meioc Dmrtb1 Esx1 Meiob Prm2 Tnp2
SSC marker genes
Progenitor genes
Diff genes
Pre-meiotic genes
Average Expression
Percent Expressed
F3
Diff2
Diff1
Progenitor
SSC5
SSC4
SSC3
SSC2
SSC1
Id4 Plvap Bcl6b Gfra1 Ret Etv5 Etv4 Lhx1 Erbb3 Tspan8 Lin28a Upp1 Sox3 Nanos3 Neurog3 Rarg Stra8 Kit Sohlh1 Sycp3 Meioc Dmrtb1 Esx1 Meiob Prm2 Tnp2
SSC marker genes
Progenitor genes
Diff genes
Pre-meiotic genes
Average Expression
Percent Expressed

**Supplementary Figure 5**. Identification of THY1$^{+}$ testicular cells in the F2 and F3 generation. (a) UMAP plots showing the nine major cell types identified from F2 and F3 PND6 testes. (b) UMAP visualization of cell distribution in each treatment group. (c) Dot plot depicts the scaled average expression of selected marker genes within each cluster.

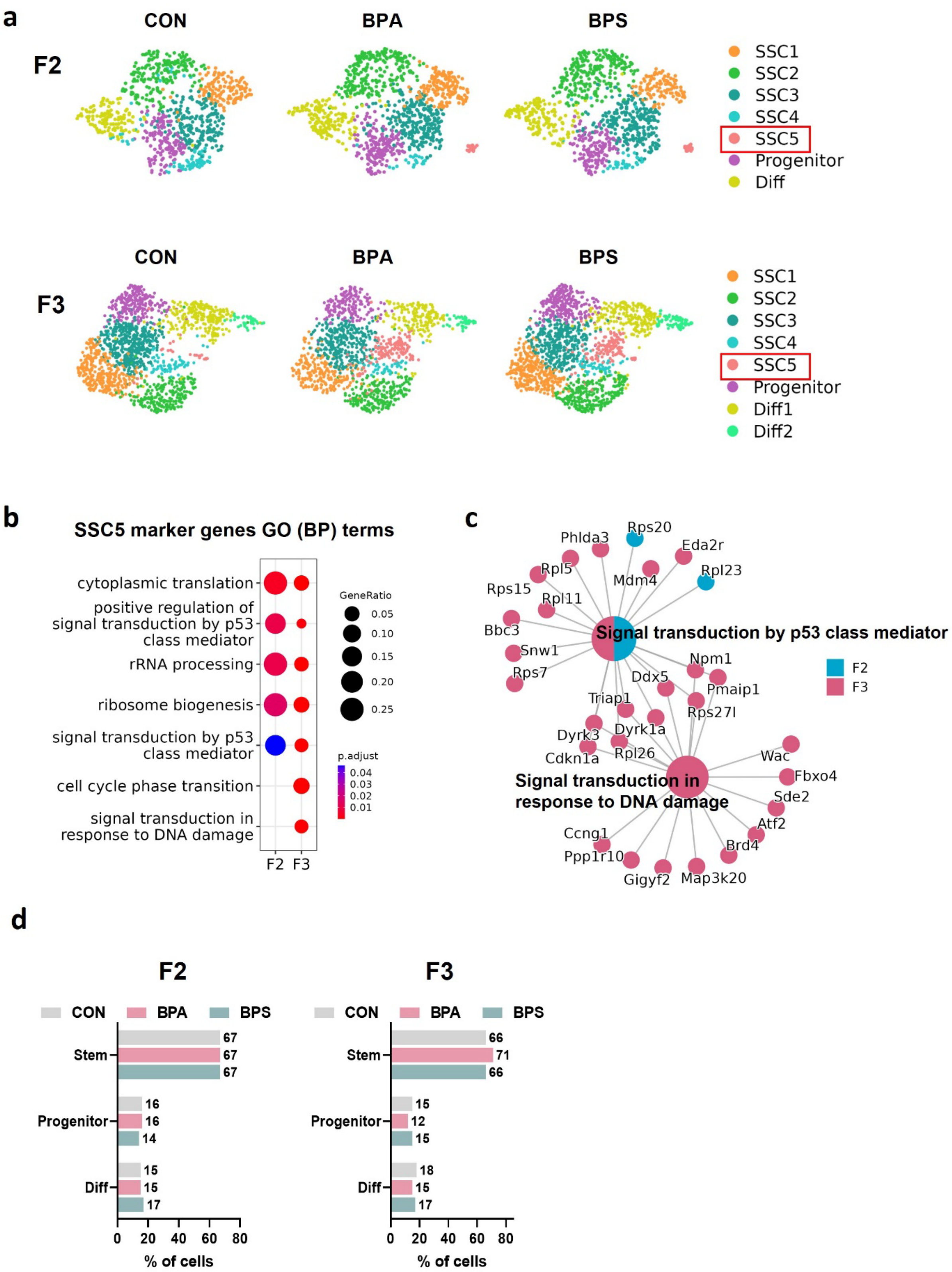
a
CON
BPA
BPS
F2
SSC1
SSC2
SSC3
SSC4
SSC5
Progenitor
Diff
F3
SSC1
SSC2
SSC3
SSC4
SSC5
Progenitor
Diff1
Diff2
b
SSC5 marker genes GO (BP) terms
cytoplasmic translation
positive regulation of signal transduction by p53 class mediator
rRNA processing
ribosome biogenesis
signal transduction by p53 class mediator
cell cycle phase transition
signal transduction in response to DNA damage
F2 F3
GeneRatio
0.05
0.10
0.15
0.20
0.25
p.adjust
0.04
0.03
0.02
0.01
c
Signal transduction by p53 class mediator
Signal transduction in response to DNA damage
F2
F3
Rps20
Phlda3
Eda2r
Rpl5
Rpl23
Mdm4
Rps15
Rpl11
Bbc3
Snw1
Rps7
Npm1
Ddx5
Pmaip1
Triap1
Rps27l
Dyrk1a
Dyrk3
Cdkn1a
Rpl26
Wac
Fbxo4
Sde2
Atf2
Ccng1
Brd4
Ppp1r10
Gigyf2
Map3k20
d
F2
F3
CON
BPA
BPS
Stem
Progenitor
Diff
67
67
67
16
16
14
15
15
17
66
71
66
15
12
15
18
15
17
0
20
40
60
80
% of cells

**Supplementary Figure 6**. Characterization of the F2 and F3 germ cell composition. (a) UMAP visualization of the distribution of germ cell subclusters in the control (CON), BPA, and BPS exposure groups. (b) GO annotation of F2 and F3 SSC5 marker genes. (c) Gene-concept network (cnetplot) of F2 and F3 SSC5 markers associated with GO terms of “Signal transduction by p53 class mediator” and “Signal transduction response to DNA damage”. SSC, spermatogonial stem cell. (d) The proportions of the SSCs, progenitors, and differentiating cells between groups in the F2 and F3 generations.

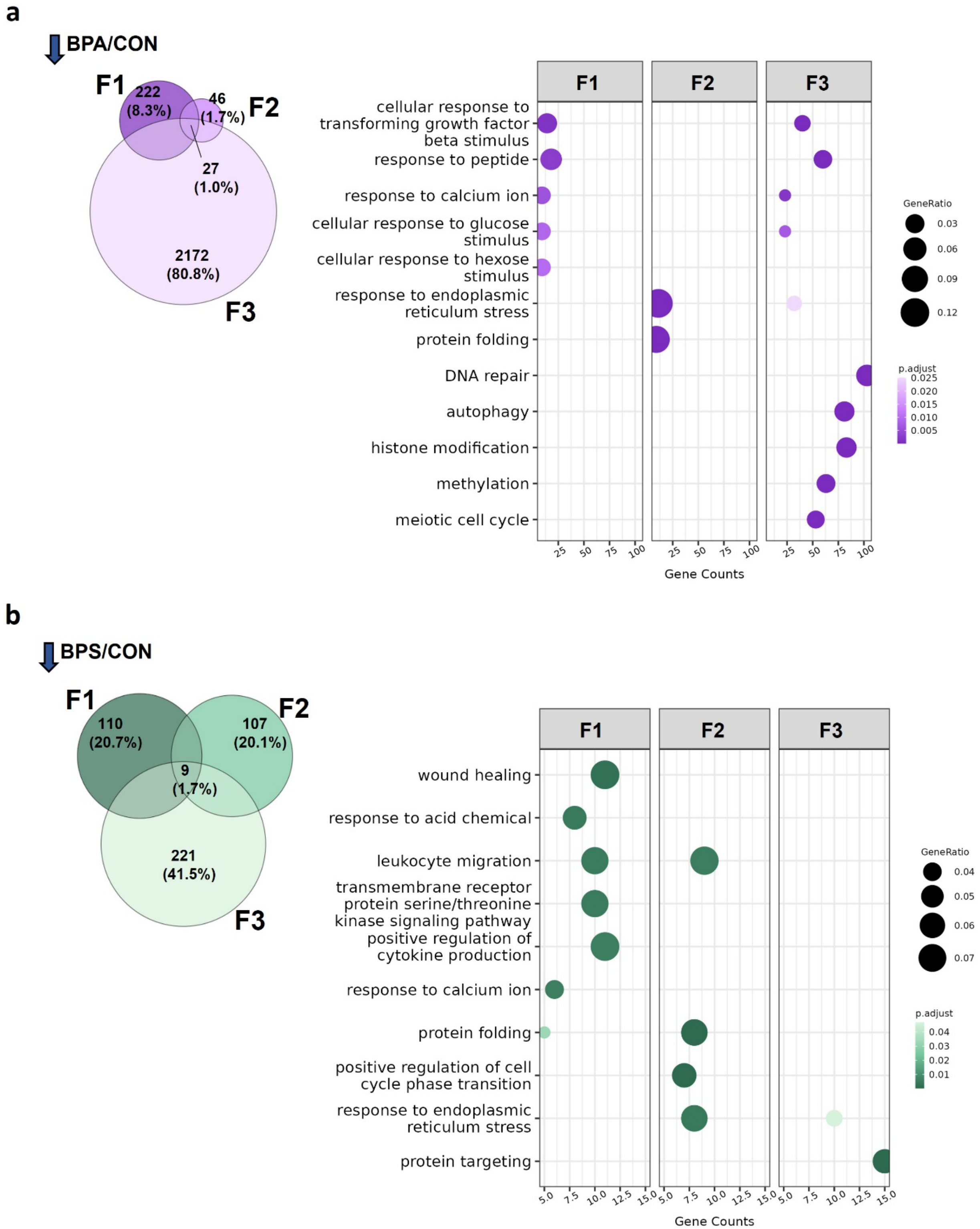
a
BPA/CON
F1
222
(8.3%)
46
(1.7%)
F2
27
(1.0%)
2172
(80.8%)
F3
F1
F2
F3
cellular response to transforming growth factor beta stimulus
response to peptide
response to calcium ion
cellular response to glucose stimulus
cellular response to hexose stimulus
response to endoplasmic reticulum stress
protein folding
DNA repair
autophagy
histone modification
methylation
meiotic cell cycle
25 50 75 100
Gene Counts
GeneRatio
0.03
0.06
0.09
0.12
p.adjust
0.025
0.020
0.015
0.010
0.005
b
BPS/CON
F1
110
(20.7%)
F2
107
(20.1%)
9
(1.7%)
221
(41.5%)
F3
F1
F2
F3
wound healing
response to acid chemical
leukocyte migration
transmembrane receptor protein serine/threonine kinase signaling pathway
positive regulation of cytokine production
response to calcium ion
protein folding
positive regulation of cell cycle phase transition
response to endoplasmic reticulum stress
protein targeting
5.0 7.5 10.0 12.5 15.0
Gene Counts
GeneRatio
0.04
0.05
0.06
0.07
p.adjust
0.04
0.03
0.02
0.01

**Supplementary Figure 7**. Transgenerational analysis of (a) BPA and (b) BPS exposure down-regulated genes in germ cells. Down-regulated genes and enriched GO terms in F1, F2, and F3 germ cells exposed to BPA or BPS.
